# Development of a cell-based DIFF-rGFP assay system for generalized discovery of viral protease Inhibitors

**DOI:** 10.1101/2024.02.01.577684

**Authors:** Wu Xiaoyan, Feng Yan, Chen Ruiting, Fang Chenjie, Wang Shuai, Sun huimin, Jian shuling, Song jiiasheng, Wu Beibei

## Abstract

Viral protease is an attractive target for antiviral therapeutics, but current viral protease inhibitor screening methods still need to be improved. Here, we systematically investigated the sites that may accommodate exogenous short peptides within Enhanced Green Fluorescent Protein (EGFP) and constructed a series of recombinant green fluorescent proteins (rGFPs). Meanwhile, a cell-based, simple and reliable assay system named DIFF-rGFP was developed relying on the co-expression of rGFP and the protease for protease inhibitor screening with the example of 3CLpro, in which the fluorescence intensity increases with the action of the inhibitor. The DIFF-rGFP assay avoided the requirement of a higher biosafety lab and can be performed in a high-throughput manner. For proof of concept, we demonstrated this method to discover novel inhibitors against SARS-CoV-2. We believe the proposed method, in combination with available drug libraries, may accelerate the identification of novel antivirals.

## 2 Introduction

Many viruses, including Coronavirus, express their proteins as a polyprotein containing one or more proteases. Such proteases autocatalytically release themselves from the precursor and cleave the remaining parts of the polyprotein into functional proteins[1] ^1^. Some medically essential viruses–including retroviruses, flaviviruses, coronaviruses, and herpesviruses–code for a protease, which is indispensable for viral maturation and pathogenesis. Inhibition of this proteolytic activity can block the production of infectious viral progeny and reduce pathogenic processes. Viral protease inhibitors have become a critical targets in developing novel antiviral drugs^2^.

SARS-CoV-2 3CLpro is a non-structural protein that cleaves at least 11 sites on large viral polyproteins essential for replication and transcription^3;4^. Therefore, inhibiting the activity of 3CLpro could disrupt the SARS-CoV-2 viral replication, making it an attractive target for COVID-19 treatment. Previous independent research groups have focused on 3CLpro, leveraging repurposed antiviral drugs such as Ribavirin, Remdesivir, Favipiravir, and Lopinavir, which are originally designed to impede viral replication. Furthermore, the structure of 3CLpro is highly conserved across Coronaviruses, indicating it may be possible to develop pan-Coronavirus inhibitors^5^.

Most frequently, the development of antiviral agents has been greatly hampered by the lack of easily controlled, cell-based assays that can be performed under biosafety level 2 (BSL2), hindering a broad application in various labs with an easy-to-prepared mode. Several existing experimental methods have been used to identify protease inhibitors. For example, some researchers have used purified proteins to screen the SARS-CoV-2 3CLpro protein *in vitro*^6789101112^. However, these experiments at the non-cellular level cannot explain whether the candidate drug will be affected by cell permeability and metabolism. In addition, inhibitors can be identified by directly assessing virus replication in cell culture, but this requires a biosafety level 3 (BSL3) condition, which may limit their high-throughput capabilities. Also in these assays, viral replication is usually quantified by plaque or quantitative RT-PCR, which may lead to difficulties distinguishing antiviral compounds from cytotoxic activity, for cytotoxicity results in reduced viral yield as well. Recently, a cell-based BSL-2 reporter system was established to identify SARS-CoV-2 3CLpro inhibitors^13^. However, based on a fluorescent reporter system called Flip-GFP, this system represents a loss-of-function approach that may lead to difficult discrimination of cytotoxicity by 3CLpro inhibition ^14^. Finally, virtual screening and molecular dynamics simulations^15^ have been widely used to identify candidate inhibitors for experimental follow-up.

Here, we describe a novel fluorescence-based report assay system to identify viral protease inhibitors in living cells. Compared to live virus screening method, our assay system is safer and simpler. The effect of inhibitor could be evaluated directly with the naked eyes. Also, the system allows for antiviral drug screening with high-throughput compatible sample processing and analysis.

## 3. Materials and Methods

### 3.1 Plasmids, cells, viruses, and drug candidate

**s** pcDNA3.1(+) expression vector was preserved by our laboratory. 293T human embryonic kidney cells were obtained from the American Type Culture Collection (Manassas, VA, USA; catalogue #CRL-3216) and maintained in Dulbecco’s modified Eagle’s medium supplemented with 2% fetal bovine serum (FBS). SARS-CoV-2 (wild type and BA. 5.2 strain) was provided by the Center for Disease Control and Prevention in Zhejiang province, China. GC376 was obtained from Aladdin. Ensitrelvir and Lopinavir were obtained from MedChemExpress.

### 3.2 Plasmid Construction

Recombinant EGFP (rGFP) was designed by inserting the protease cleavage sequence into the EGFP coding sequence. The insertion sites were summarized in **Table 1**. PCR amplification was performed by Prime Star DNA polymerase (Takara). The GFP and rGFP were cloned into the multiple cloning site (MCS) of the pcDNA3.1(+) vector followed by the Uniclone One Step Seamless Cloning Kit instruction, respectively, producing the vector of pGFP and a series of prGFPs. Subsequently, the viral protease was cloned at the 3’terminal of the rGFP linked by a 2A peptide sequence, which produced plasmids co-expressing rGFP and SARS-CoV-2 3CLpro named prGFP-3CLpro. The protease cleavage sequences are summarized in **Table 2**.

**Table 1.** A summary of the insertion sites 1∼45.

| Name | Location | Name | Location |
| --- | --- | --- | --- |
| Site1 | W58-P59 | Site24 | K159-N160 |
| Site2 | N145-Y146 | Site25 | T10-G11 |
| Site3 | H149-N150 | Site26 | Q70-C71 |
| Site4 | D77-H78 | Site27 | N160-G161 |
| Site5 | F84-F85 | Site28 | G175-S176 |
| Site6 | Y93-V94 | Site29 | Q205-S206 |
| Site7 | K102-D103 | Site30 | V220-L221 |
| Site8 | T119-L120 | Site31 | D118-T119 |
| Site9 | N106-Y107 | Site32 | L120-V121 |
| Site10 | E133-D134 | Site33 | V121-N122 |
| Site11 | E173-D174 | Site34 | A155-D156 |
| Site12 | K210-D211 | Site35 | D156-K157 |
| Site13 | K215-R216 | Site36 | K157-Q158 |
| Site14 | P197-D198 | Site37 | N171-I172 |
| Site15 | Q158-K159 | Site38 | I172-E173 |
| Site16 | G25-H26 | Site39 | D174-G175 |
| Site17 | Y40-G41 | Site40 | S176-V177 |
| Site18 | P55-V56 | Site41 | V177-Q178 |
| Site19 | G139-H140 | Site42 | I153-M154 |
| Site20 | G117-D118 | Site43 | M154-A155 |
| Site21 | P193-V194 | Site44 | R169-H170 |
| Site22 | F115-E116 | Site45 | H170-N171 |
| Site23 | E116-G117 |  |  |

**Table 2.**
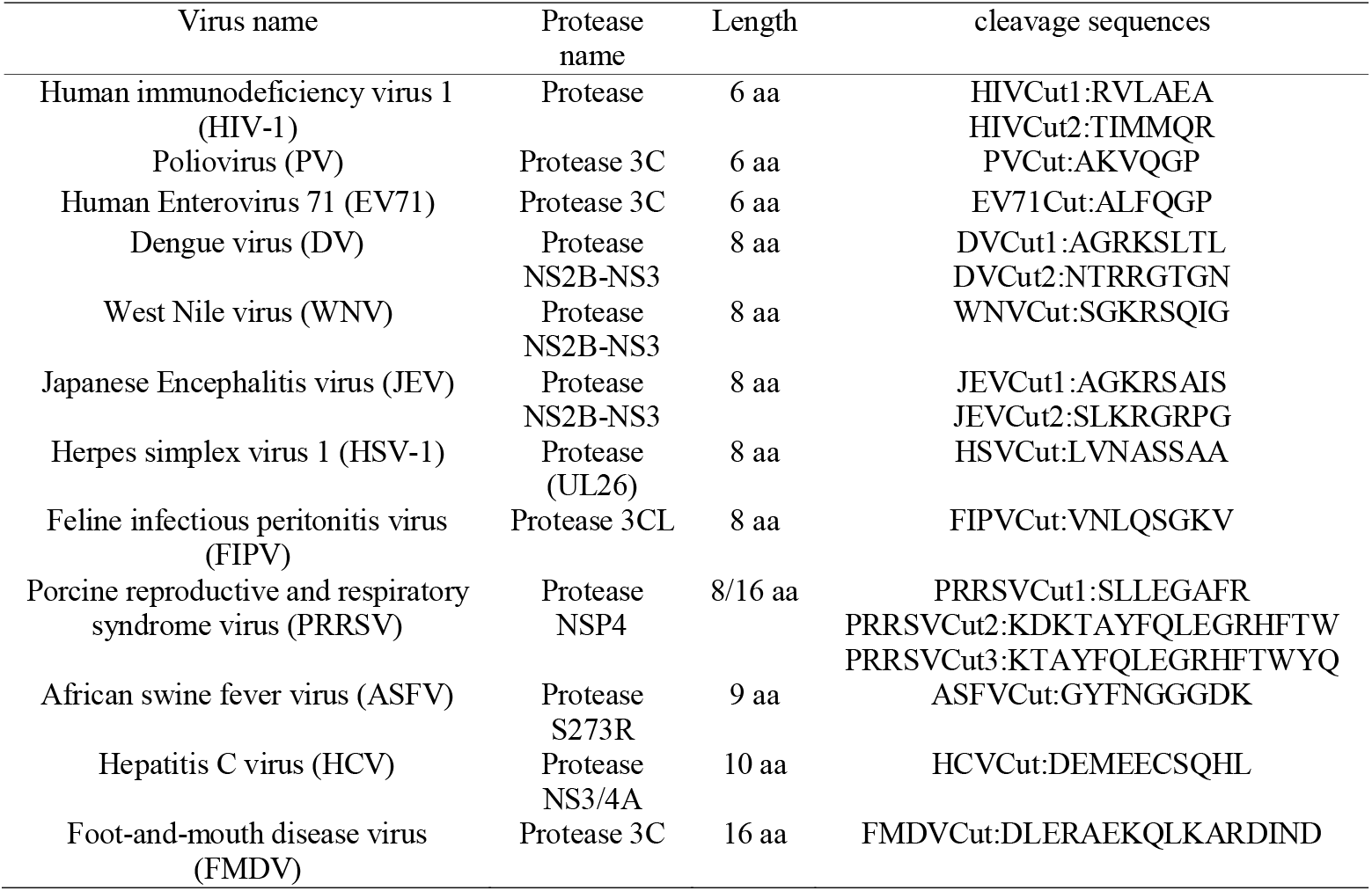
The viral cleavage sequences.

### 3.3 Drug candidates screening and rGFP assay development

Prepare 293T cells with a cell density of 5X10^5^ cells/well for a 24-well plate. Plasmid transfections were carried out using lipofectamine 2000 according to the instructions. The drug candidates were dissolved in DMSO and diluted with DMEM. 4-6 hrs after transfection, the different concentrations of drug candidates were added into the corresponding wells and culture for another 48h. The green fluorescence was detected by a fluorescent microscope (OLYMPUS, CKX53).

### 3.4 Assay Validation by High-throughput Screening

Before the experiment, the Maximum non-toxic concentrations of small molecular compounds on Vero-E6 cells were investigated using Cell Counting Kit-8 (CCK-8) (Beyotime, Shanghai, China). The concentration with >90% cell survival rate was the maximum non-toxic concentration.

For the live virus experiment, Vero-E6 cells seeded in a 96-well plate were infected with 0.01 MOI SARS-CoV-2 (wild type or Omicron strain) at 37L for 1 hr. After infection, the virus was discarded and the cells were treated with 100 μL different concentrations of drug candidates (6.25-0.39 μM for GC376, and 3.125-0.19 μM for Ensitrelvir). For the virus control (VC) group, cells were covered with 100 μL/well MM after infection. Cell control refers to cells without infection. After incubation at 37L for 3 days, the cells were rinsed with PBS twice and stained with MEM containing 10% CCK-8 for 1.5 hr. The optical density values of each well were measured at 450 nm, and the cell viability was calculated.

For rGFP assay, the 293T cells were seeded on a 96-well plate at 2.5X104 cells/well, the co-expression plasmid was transfected following the lipofectamine 2000 transfection reagent kit instruction when the cell convergence reached 70-80%. Replace the culture medium with fresh medium added with different concentrations of drug candidates 6hrs later and culture the cells for 2 days at 37□, 5% CO2. When detecting, the medium was discarded and the fluorescent intensity was read by multifunctional Enzyme Labeling Instrument at 485nm excitation light. Positive control and Negative control were set as the cells transfected with GFP-expressing plasmid and cells transfected with pcDNA 3.1(+), respectively.

### 3.5 Statistical Analysis

Statistical analysis was performed in GraphPad Prism 9.2.0 (GraphPad Software, San Diego, CA, USA). In figure 6, EC50 value was obtained by nonlinear regression that was performed for each data set with the best-fit value. For all experiments, at least three independent biological replicates were performed.

## 4. Results

### 4.1 Screening of SARS-CoV-2 3CLpro cleavage sequences insertion site that do not affect GFP function

We totally selected 45 sites for SARS-CoV-2 3CLpro protease cleavage sequence compatibility screening. It can be observed that among the selected 45 insertion sites, only 19 sites, i.e. site 8,11,15,19,20,23,24,25,28,31,34,35,36,37,38,39,40,43,45 (note: “/” indicates “or,” and the same applies throughout), are compatible with the SARS-CoV-2 3CLpro protease cleavage sequence without significantly affecting the fluorescence of GFP or with only limited impact. At the remaining sites, such as site6, site10, site 13, site 22 and 27, the fluorescence of GFP cannot be detected after inserting the 3CLpro protease cleavage sequence (Fig.1).

**Fig. 1.**
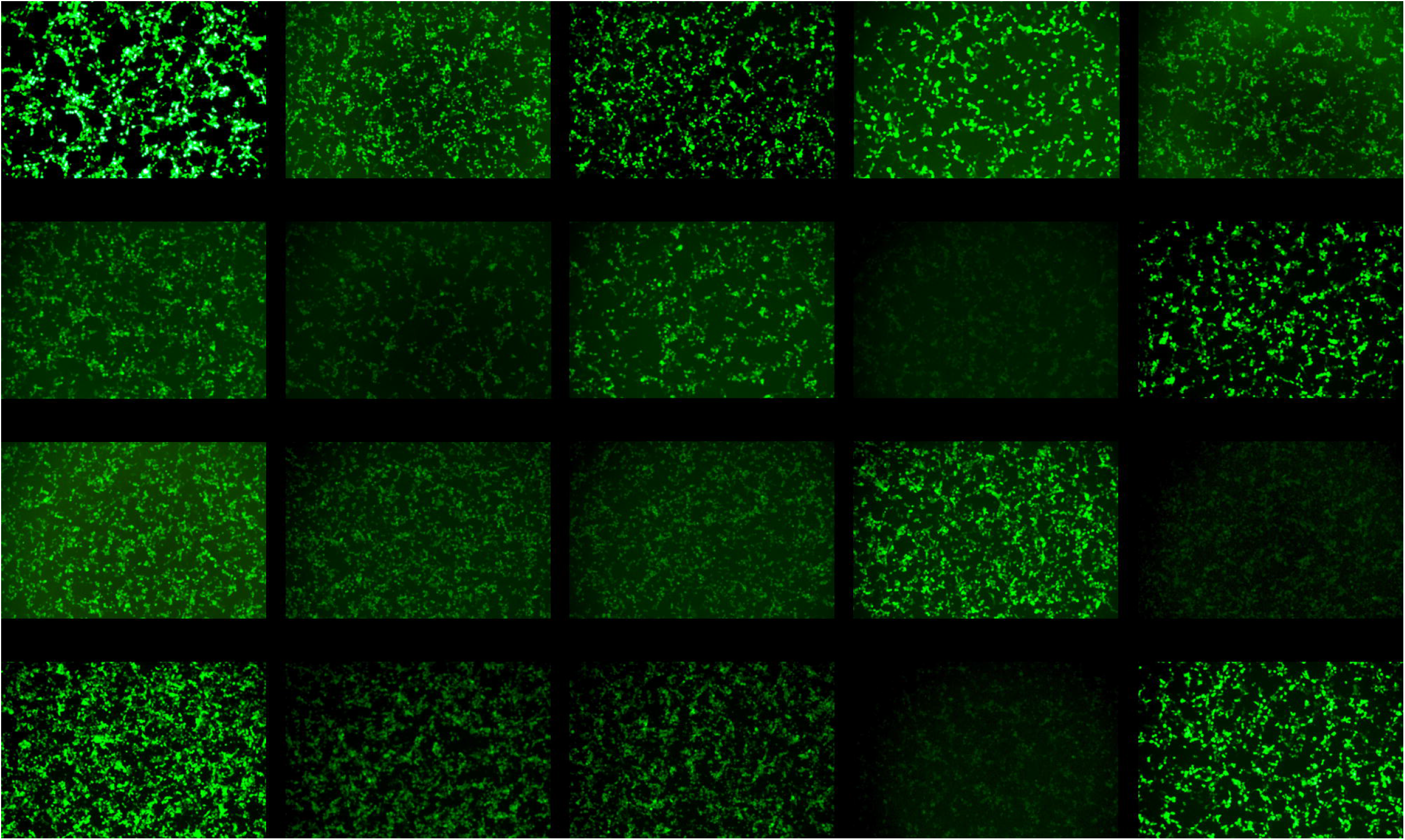
Transfection of the plasmid expressing recombinant green fluorescent proteins (rGFPs) into 293T cells. rGFP1∼45 represent recombinant fluorescent proteins with insertion of SARS-CoV-2 3CLpro cleavage sequences into the corresponding site1∼45 in Table1. For instance, rGFP8 represents the insertion of the 3CLpro cleavage sequence between T119 and L120 (site8) of GFP.

By analyzing the above sites, three highly flexible regions that may accommodate exogenous short peptides are found. Region 1:E116∼L120 (Table S1), the amino acid sequence of the above region is EGDTL, and fluorescence is not detected in the region’s near-neighboring site22, site32 and 33. Region 2: M154∼N160 (Table S2), the amino acid sequence of the above region is MADKQKN, and fluorescence is not detected in the region’s near-neighboring site42, site43 and 27. Region 3: H170∼V177 (Table S3), the amino acid sequence of the above region is HNIEDGSV, and fluorescence is not detected in the region’s near-neighboring site44 and site41.

### 4.2 Flexible regions are compatible with different or two combined cleavage sequences

Some single-stranded RNA viruses and some DNA viruses’ protease cleavage sequences were chosen for insertion into the possible flexible region site. The viral proteases and their corresponding cleavage sequence are listed in Table 2. Firstly, we selected site 15, 20 and site 45, which located in the three regions, respectively. Every viral cleavage sequence could get a positive result within these three sites but the HCV cleavage sequence. We then tried site24 and got the functional GFP (Fig.2a). The results demonstrated that for one single cleavage sequence, at least one site within the three regions would be available for insertion. Therefore, it is assumed that there are three flexible regions within the GFP which can accept exogenous cleavage sequences.

**Fig. 2.**
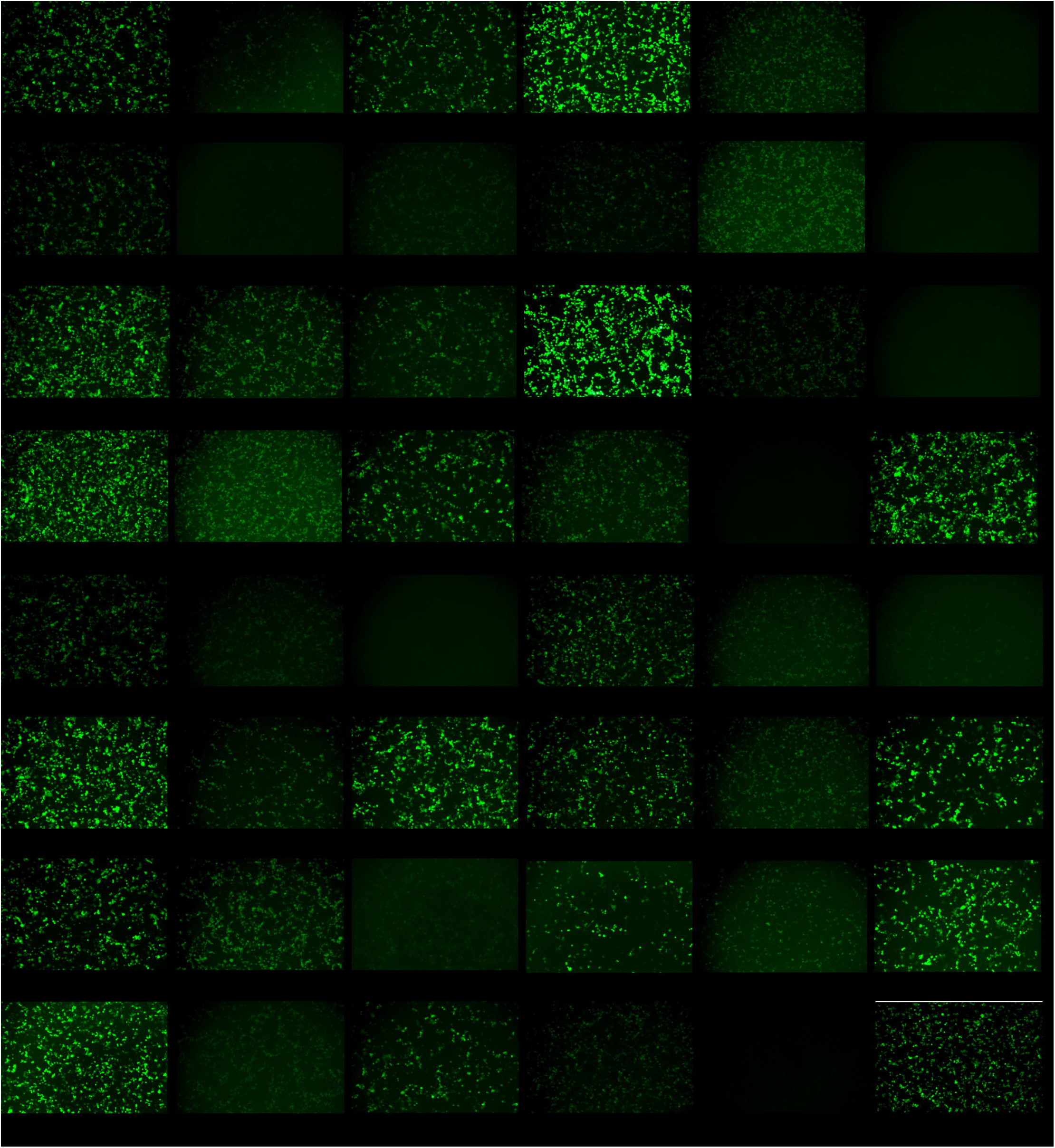
Flexible region test. **a**, Insertion of various cleavage sequences into representative sites on three regions. rGFP15-HIVCut1 represents inserting of HIVCut1 into Q158-K159 of GFP, and so on. **b**, the corresponding locations of the available sites and flexible region within GFP 3D structure(pdb:1bfp)^16^, yellow marked. The cyan represents sites that could not accommodate cleavage sequences.

**Fig. 3.**
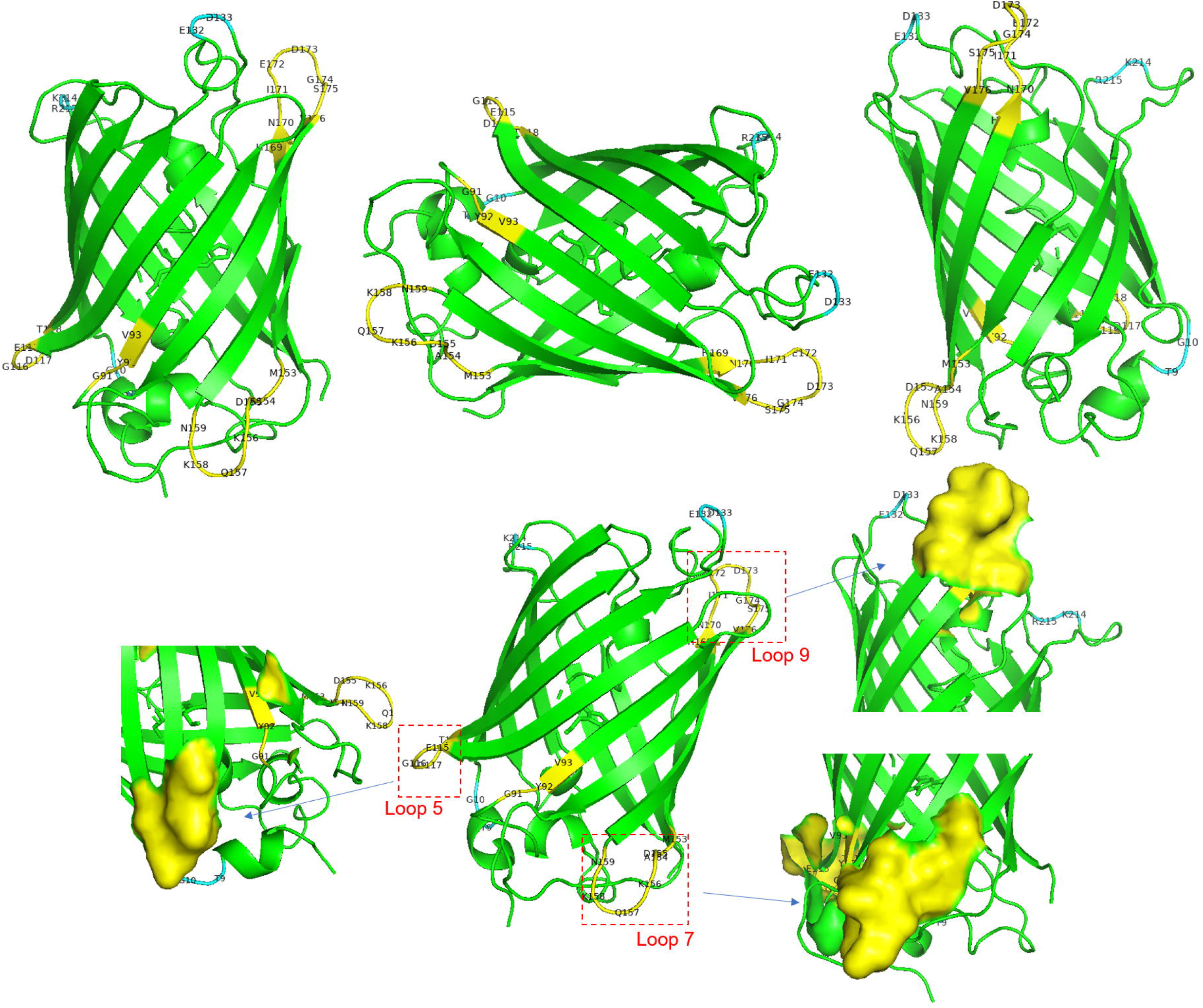
Co-expression of rGFP and protease in 293T cells.

Simultaneously, the sites and regions mentioned above were marked in GFP 3D structure (Fig2b). We could see the three regions were mainly located in the loop domain or near the loop between the beta strands of β4 to β5, β6 to β7, and β8 to β9. However, not all loops are feasible for insertion. Apart from this, we found the Y92-G91 site was not on the surface of the GFP, this may be why the lead cleavage sequence was wrapped up internally, which protease cannot digest efficiently.

### 4.3 Co-expression Viral protease with rGFP would prevent the fluorescent function of GFP

To confirm that the rGFP could be digested by the viral protease, we co-expression the rGFP with the SARS-CoV-2 3CL protease as a model. The sequence of 3CLpro was added to the rGFP expression vector with the connection of P2A. From the results of Fig. 5, no green fluorescence was detected when rGFP8, rGFP20, rGFP24, rGFP31 were co-expressed with 3CLpro protease. While, weak green fluorescence was still observed when rGFP11, rGFP15, rGFP28, rGFP36, rGFP38, rGFP39, rGFP40 and rGFP45 were co-expressed with 3CLpro protease. This may be attributed to the spatial conformation of the protease cleavage sequence, making it less favorable for the exposure and recognition of the protease cleavage sequence by 3CLpro for efficient cleavage. From then on, we confirmed that the viral protease could digest the cleavage sequence with relatively high efficiency, and it was realized that this system could be used for screening the protease inhibitors or evaluating the inhibitor. Take the SARS-CoV-2 as an example, prGFP8-3CLpro, prGFP15-3CLpro and prGFP20-3CLpro plasmids were selected for further study.

### 4.4 Development of a DIFF-rGFP assay for SARS-Cov-2 3CLpro inhibitor screening

SARS-CoV-2 polyproteins are processed by two viral proteases, papain-like protease (PLpro) and 3CLpro, which are excellent targets for the development of therapeutic antivirals. Some compounds have been validated targeting the 3CLpro, such as GC376 and Enstrelvir, but some are still needed to be investigated. Here, three compounds were chosen to develop the rGFP assay for 3CLpro inhibitor screening. With the addition of GC376 up to 10μM, the green fluorescence reoccurred during the observation time. As the concentration increased, the fluorescence became brighter (Fig.4).

**Fig. 4.**
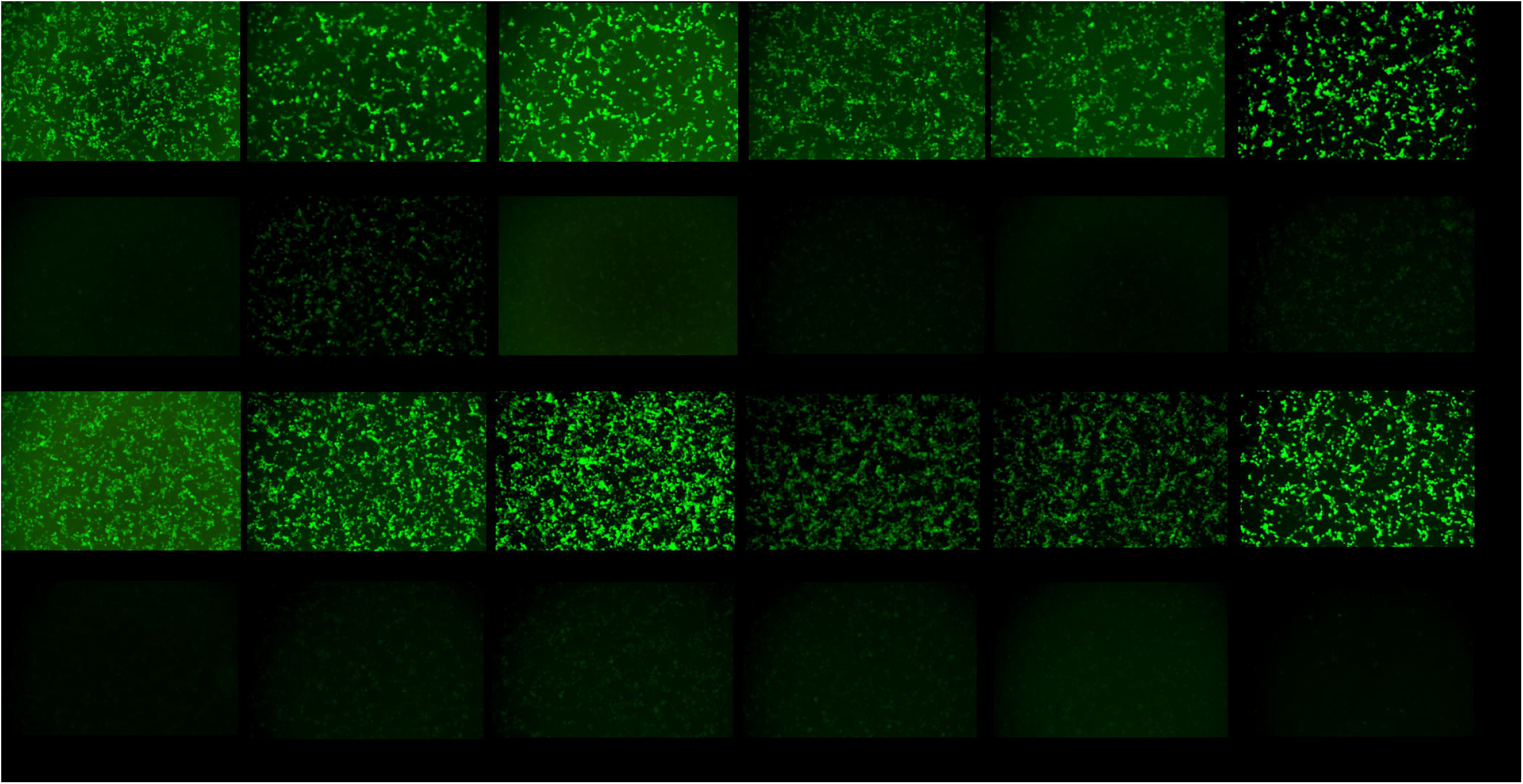
Validation of the rGFP assay of prGFP15-3CLpro. The fluorescence intensity was investigated after transfection 24h, 48h and 72h.

**Fig. 5.**
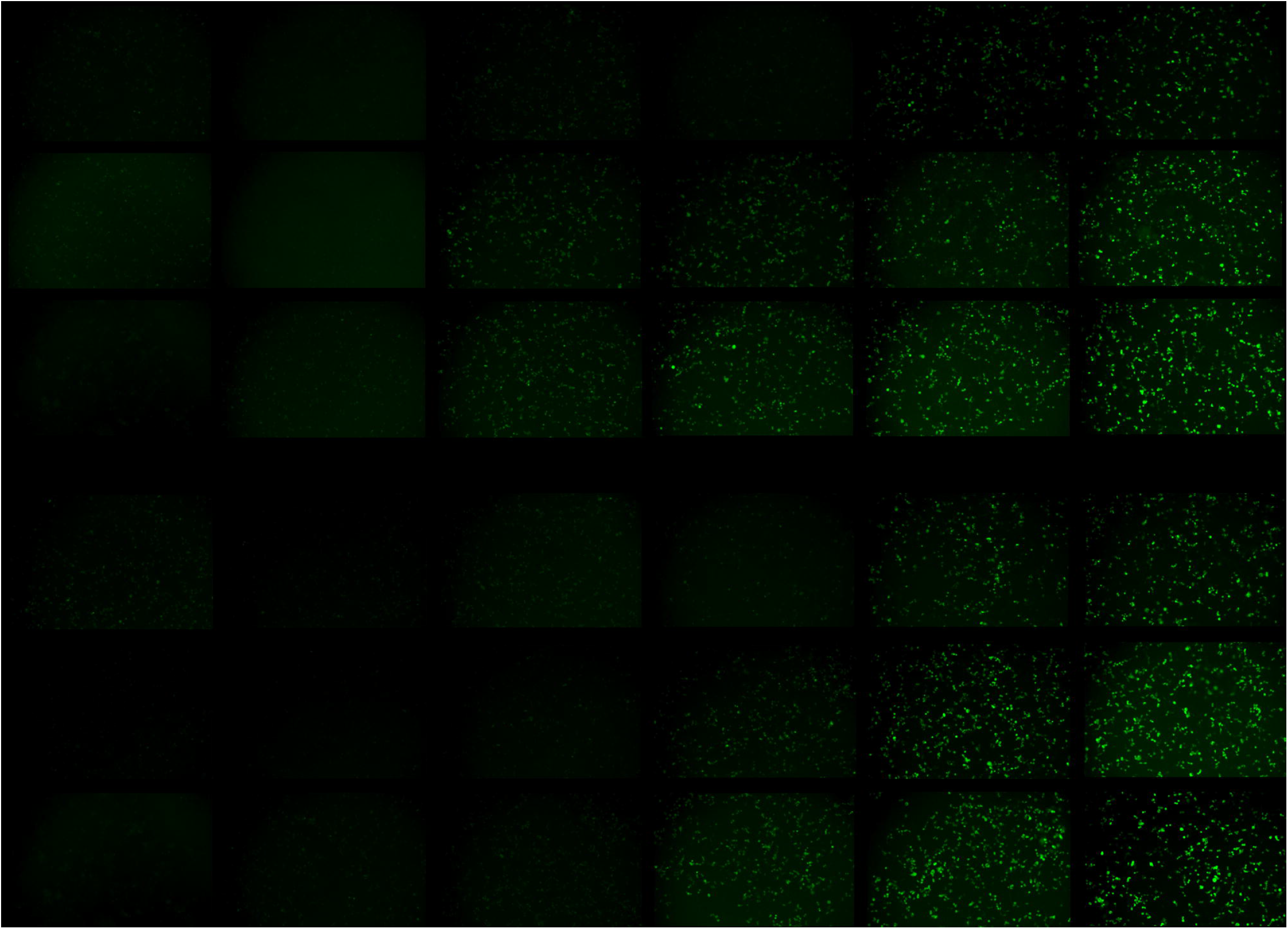
EC50 of the live virus and rGFP15-3CLpro assay.

The same result was obtained for Enstrelvir except for the difference in minimum active concentration which is 1 μM. While, lopinavir didn’t seem to work. In this stage, we compared prGFP8-3CLpro (data not shown), prGFP15-3CLpro and prGFP20-3CLpro (data not shown) and found prGFP15-3CLpro possessed the best performance for assay development.

### 4.5 Comparison of the effect of report assay system and live virus screening inhibitors

The above results have demonstrated that the prGFP15-3CLpro assay can be used to screen 3CLpro protease inhibitor candidates. To understand the differences between the prGFP15-3CLpro assay system and the live virus screening, we detected and figured out the EC50 of two small molecules, GC376 and enstrelvir, based on the cell viability after SARS-Cov-2 infection and the GFP intensity, respectively. For GC376, the EC50 was ∼6.5 μM of prGFP15-3CLpro assay, one-fold higher than that of live virus screening. For enstrelvir, the EC50 was ∼1.493 μM of prGFP15-3CLpro assay. The differences between these two kinds of assay were thought to come mainly from the reported organism. Once the drug worked, the virus replication was restrained. Otherwise, the virus proliferated exponentially, so the signal was high. However, our DIFF-rGFP assay was smoother and more moderate than virus proliferation, therefore the change in data response was not so substantial. From then on, A cell-based, simple and reliable DIFF-rGFP assay system was developed. Also, the fluorescence can be detected under a spectrometer with high-throughput in a 96-well or 384-well plate.

## 5 Discussion

GFP is one of the most commonly used fluorescent proteins in scientific research, which is usually fused with the gene of interest at its N-terminal or C-terminal without influencing the green fluorescence detection. The fluorescence expression is strictly related to the barrel-like structure, which is composed of the helical chromophore segment and eight β-strands connected by 10 solvent-accessible loops, thus, little research inserted exogenous sequences into the internal of the GFP residuals. Although one research reported that the solvent-accessible loops are candidate sites for inserting random peptides, the sites or flexible region within GFP compatible with exogenous protease cleavage sequences still lack experimental evidence^17^. Only three sites have been validated upon the HCV NS3/4A cleavage sequence ^18^.

Some viruses express their proteins as a polyprotein containing one or more proteases. Such proteases autocatalytically release themselves from the precursor and cleave the remaining parts of the polyprotein into functional proteins. Inhibition of this proteolytic activity can block the production of infectious viral progeny and reduce pathogenic processes^19,20,21,22^, these proteases inevitably become essential targets for antiviral drug development. However, efficient and accurate methods for antiviral candidates screening of protease inhibitor are still lacking. Method of using expressed protease mixed with the drug candidate, such as fluorescence resonance energy transfer(FRET)-based screening assay, reacts the inhibitory activity in vitro, but could not imitate the processing of the drug candidate entrance into the cells is short of accuracy.^13,23^ And using the live viruses to detect the candidate activity usually needs BSL-2 or above laboratory situation^13,24^. In this work, we developed an easy-going detecting platform for protease inhibitor screening, which can be manipulated in a normal bio-safety level lab. For the first time, we found the sites and flexible region competent in loading exogenous viral protease cleavage sequence without affecting or with little affection on the green fluorescence intensity compared with wt GFP based on SARS-Cov-2 3Cpro cleavage sequence and other viral protease cleavage sequence. The most significant advantage is that for the unique exogenous sequence insertion, we can certainly seek out one or more proper sites within the flexible region of GFP and the chromophore structure will not be prevented. When the exogenous sequences are inserted into the GFP, a recombinant green fluorescent reporting protein (rGFP) could be produced. Thus, researchers no longer need to take a lot of time studying from the beginning when designing rGFP or other fluorescent report proteins homologous with GFP.

Secondly, we co-expressed the viral protease and the fluorescent report protein in one plasmid. Because the two genes were promoted by a shared promoter linked by a 2A self-cleaving sequence, the plasmids added into the cell medium tend to be more conveniently controlled compared with two separated plasmids, and the expression of the protease and rGFP are more stable in different cell types. Our following research indicated that the two genes could also linked by a flexible linker and the protease cleavage sequence (Data not shown). In this stage, the key point is that the protease cuts the rGFP efficiently; otherwise, the fluorescence still exists. In some cases, we found the protease couldn’t digest the cleavage sequence, so that the system didn’t work so well. Based on the above platform, a series antiviral candidate screening assays can be developed targeting many positive single-stranded RNA viruses and some DNA viruses^25^.

Here, we also developed the rGFP reporter assays(Fig.6) for SARS-CoV-2 3CLpro and confirmed the inhibition activity of the small molecule GC376^26^ and enstrelvir^27^. The results showed that prGFP15-3CLpro was the most sensitive assay. Meanwhile in the assays, the EC50 was higher than the live virus screening. It was thought that the difference mainly came from the different characteristics between virus and plasmid. Furthermore, the assay can be utilized in high-throughput screening candidate drugs and extend to other types of viral protease.

**Fig. 6.**
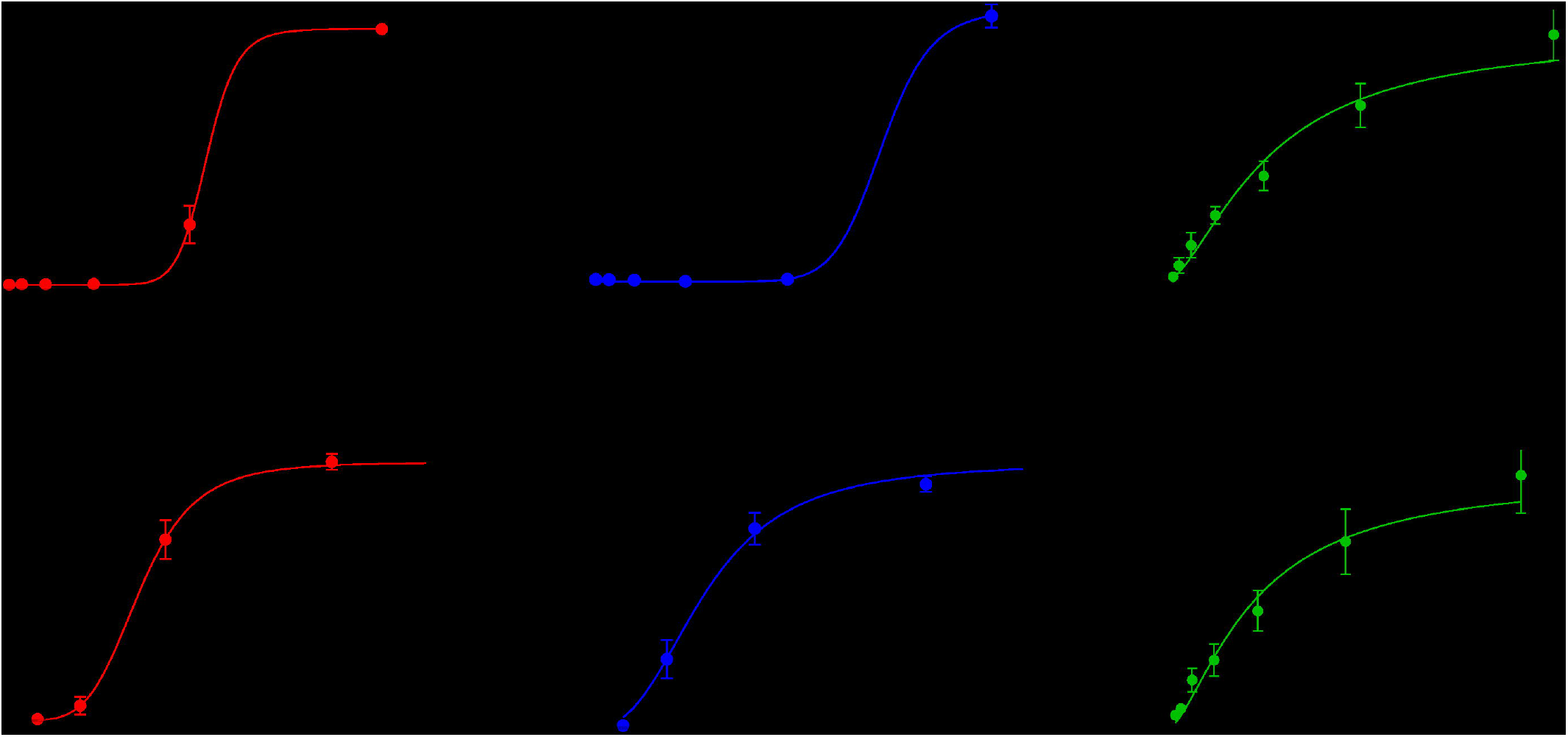
Schematic diagram of rGFP reporter assay.

In summary, our study thoroughly investigated compatible sites or regions within the GFP and provided a lab-friendly, accurate, efficient way for viral protease inhibitors screening, offering a framework to accelerate the time and effort in a cost-effective manner.

---

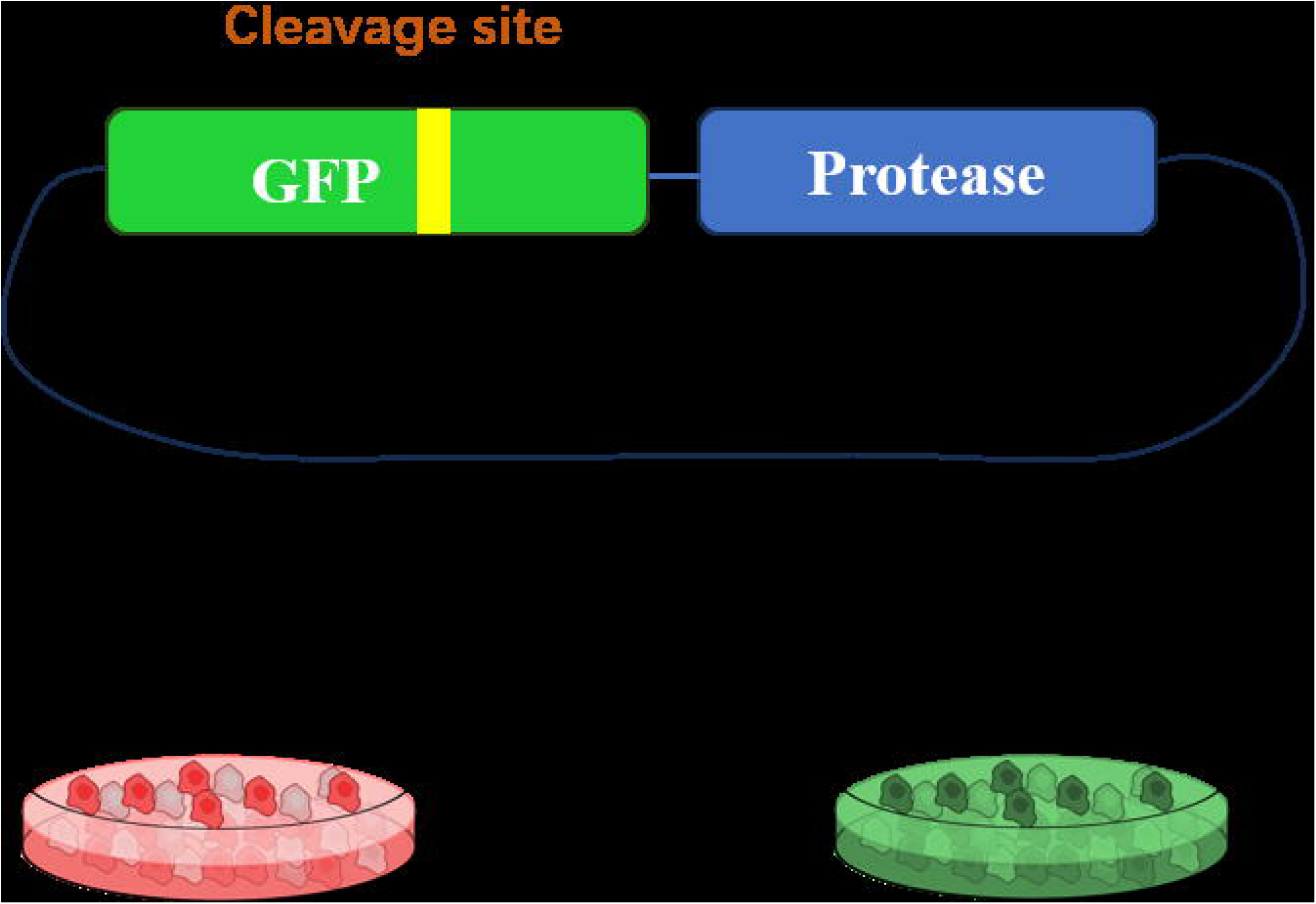

## References

1. Tsu, B.V., Fay, E.J., Nguyen, K.T., Corley, M.R., Hosuru, B., Dominguez, V.A., Daugherty, M.D., 2021. Running with scissors: evolutionary conflicts between viral proteases and the host immune system. Front. Immunol. 12, 769543

2. Majerová T, Konvalinka J. Viral proteases as therapeutic targets. Mol Aspects Med. 2022 Dec;88:101159. doi: 10.1016/j.mam.2022.101159. Epub 2022 Nov 29. PMID: 36459838; PMCID: PMC9706241.

3. Durai, P., Batool, M., Shah, M. & Choi, S. Middle East respiratory syndrome coronavirus: Transmission, virology and therapeutic targeting to aid in outbreak control. Exp. Mol. Med. 47, e181–e181 (2015).

4. Arya R, Kumari S, Pandey B, Mistry H, Bihani SC, Das A, Prashar V, Gupta GD, Panicker L, Kumar M. Structural insights into SARS-CoV-2 proteins. J Mol Biol. 2021 Jan 22;433(2):166725. doi: 10.1016/j.jmb.2020.11.024. Epub 2020 Nov 24. PMID: 33245961; PMCID: PMC7685130.)

5. Khan, M.T.; Ali, A.; Wang, Q.; Irfan, M.; Khan, A.; Zeb, M.T.; Zhang, Y.-J.; Chinnasamy, S.; Wei, D.Q. Marine natural compounds as potents inhibitors against the main protease of SARS-CoV-2-a molecular dynamic study. J. Biomol.Struct. Dyn. 2020, 1–11.

6. Fu, L.; Ye, F.; Feng, Y.; Yu, F.; Wang, Q.; Wu, Y.; Zhao, C.; Sun, H.; Huang, B.; Niu, P.; et al. Both Boceprevir and GC376 efficaciously inhibit SARS-CoV-2 by targeting its main protease. Nat. Commun. 2020, 11, 4417.

7. Hung, H.-C.; Ke, Y.-Y.; Huang, S.Y.; Huang, P.-N.; Kung, Y.-A.; Chang, T.-Y.; Yen, K.-J.; Peng, T.-T.; Chang, S.-E.; Huang, C.-T.; et al. Discovery of M Protease Inhibitors Encoded by SARS-CoV-2. Antimicrob. Agents Chemother. 2020, 64, e00872-20. Inhibitors by a Quantitative High-Throughput Screening. ACS Pharmacol. Transl. Sci. 2020, 3, 1008–1016.

8. Ma, C.; Sacco, M.D.; Hurst, B.; Townsend, J.A.; Hu, Y.; Szeto, T.; Zhang, X.; Tarbet, B.; Marty, M.T.; Chen, Y.; et al. Boceprevir, GC-376, and calpain inhibitors II, XII inhibit SARS-CoV-2 viral replication by targeting the viral main protease. Cell Res. 2020, 30, 678–692.

9. Jin, Z.; Du, X.; Xu, Y.; Deng, Y.; Liu, M.; Zhao, Y.; Zhang, B.; Li, X.; Zhang, L.; Peng, C.; et al. Structure of M(pro) from SARS-CoV-2 and discovery of its inhibitors. Nature 2020, 582, 289–293.

10. Rathnayake, A.D.; Zheng, J.; Kim, Y.; Perera, K.D.; Mackin, S.; Meyerholz, D.K.; Kashipathy, M.M.; Battaile, K.P.; Lovell, S.; Perlman, S.; et al. 3C-like protease inhibitors block coronavirus replication in vitro and improve survival in MERS-CoV-infected mice. Sci. Transl. Med. 2020, 12, eabc5332.

11. Wang, Y.-C.; Yang, W.-H.; Yang, C.-S.; Hou, M.-H.; Tsai, C.-L.; Chou, Y.-Z.; Hung, M.-C.; Chen, Y. Structural basis of SARS-CoV-2 main protease inhibition by a broad-spectrum anti-coronaviral drug. Am. J. Cancer. Res. 2020, 10, 2535–2545.

12. Zhu, W.; Xu, M.; Chen, C.Z.; Guo, H.; Shen, M.; Hu, X.; Shinn, P.; Klumpp-Thomas, C.; Michael, S.G.; Zheng, W. Identification of SARS-CoV-2 3CL Protease

13. Rawson JMO, Duchon A, Nikolaitchik OA, Pathak VK, Hu WS. Development of a Cell-Based Luciferase Complementation Assay for Identification of SARS-CoV-2 3CLpro Inhibitors. Viruses. 2021;13(2):173. Published 2021 Jan 24. doi:10.3390/v13020173

14. Froggatt, H.M.; Heaton, B.E.; Heaton, N.S. Development of a Fluorescence-Based, High-Throughput SARS-CoV-2 3CL(pro) Reporter Assay. J. Virol. 2020, 94, e01265–20.

15. Chan, LC., Mat Yassim, A.S., Ahmad Fuaad, A.A.H. et al. Inhibition of SARS-CoV-2 3CL protease by the anti-viral chimeric protein RetroMAD1. Sci Rep 13, 20178 (2023). 10.1038/s41598-023-47511-z

16. Wachter RM, King BA, Heim R, Kallio K, Tsien RY, Boxer SG, Remington SJ. Crystal structure and photodynamic behavior of the blue emission variant Y66H/Y145F of green fluorescent protein. Biochemistry. 1997 Aug 12;36(32):9759–65. doi: 10.1021/bi970563w. PMID: 9245407.

17. Abedi MR, Caponigro G, Kamb A. Green fluorescent protein as a scaffold for intracellular presentation of peptides. Nucleic Acids Res. 1998;26(2):623–630. doi:10.1093/nar/26.2.623

18. Yan Xing, Shi Mei-lei, Chen Na, Wang Yang, Zhang Yan, Wang Zheng, Li Xiao-chen, Zhao zhi-hu, “Construction of a Fluorescence Detection Method For HCV NS3/4Ap Protease Activity. J. China Biotechnology, 2012,32(07):84–88. doi:10.13523/j.cb.20120715.

19. Schneider, J., Kent, S.B., 1988. Enzymatic activity of a synthetic 99 residue protein corresponding to the putative HIV-1 protease. Cell 54 (3), 363–368.

20. Han, D.S., Hahm, B., Rho, H.M., Jang, S.K., 1995. Identification of the protease domain in NS3 of hepatitis C virus. J. Gen. Virol. 76 (Pt 4), 985–993.

21. Tsu, B.V., Fay, E.J., Nguyen, K.T., Corley, M.R., Hosuru, B., Dominguez, V.A., Daugherty, M.D., 2021. Running with scissors: evolutionary conflicts between viral proteases and the host immune system. Front. Immunol. 12, 769543

22. Majerová T, Konvalinka J. Viral proteases as therapeutic targets. Mol Aspects Med. 2022;88:101159. doi:10.1016/j.mam.2022.101159

23. Rothan HA, Teoh TC. Cell-Based High-Throughput Screening Protocol for Discovering Antiviral Inhibitors Against SARS-COV-2 Main Protease (3CLpro). Mol Biotechnol. 2021;63(3):240–248. doi:10.1007/s12033-021-00299-7

24. Rothan HA, Teoh TC. Cell-Based High-Throughput Screening Protocol for Discovering Antiviral Inhibitors Against SARS-COV-2 Main Protease (3CLpro). Mol Biotechnol. 2021;63(3):240–248. doi:10.1007/s12033-021-00299-7

25. Majerová T, Konvalinka J. Viral proteases as therapeutic targets. Mol Aspects Med. 2022;88:101159. doi:10.1016/j.mam.2022.101159

26. Heilmann E, Costacurta F, Moghadasi SA, et al. SARS-CoV-2 3CLpro mutations selected in a VSV-based system confer resistance to nirmatrelvir, ensitrelvir, and GC376. Sci Transl Med. 2023;15(678):eabq7360. doi:10.1126/scitranslmed.abq7360

27. McKimm-Breschkin JL, et al. COVID-19, Influenza and RSV: Surveillance-informed prevention and treatment - Meeting report from an isirv-WHO virtual conference. Antiviral Res. 2022;197:105227.

