## Supplementary material for "Development of a cell-based DIFF-rGFP assay system for generalized discovery of viral protease Inhibitors": Table S1, Table S2, Table S3

**Supplementary Materials**

Table S1. Region 1 and adjacent sites

| Site | Location | Fluorescence Detected |
| --- | --- | --- |
| Site22 | F115-E116 | NO |
| Site23 | E116-G117 | YES |
| Site20 | G117-D118 | YES |
| Site31 | D118-T119 | YES |
| Site8 | T119-L120 | YES |
| Site32 | L120-V121 | NO |

Table S2. Region 2 and adjacent sites

| Site | Location | Fluorescence Detected |
| --- | --- | --- |
| Site34 | A155-D156 | YES |
| Site35 | D156-K157 | YES |
| Site36 | K157-Q158 | YES |
| Site15 | Q158-K159 | YES |
| Site24 | K159-N160 | YES |
| Site27 | N160-G161 | NO |
| Site42 | I153-M154 | NO |
| Site43 | M154-A155 | NO |

Table S3. Region 3 and adjacent sites

| Site | Location | Fluorescence Detected |
| --- | --- | --- |
| Site44 | R169-H170 | NO |
| Site45 | H170-N171 | YES |
| Site37 | N171-I172 | YES |
| Site38 | I172-E173 | YES |
| Site11 | E173-D174 | YES |
| Site39 | D174-G175 | YES |
| Site28 | G175-S176 | YES |
| Site40 | S176-V177 | YES |
| Site41 | V177-Q178 | NO |
